# Identification and current palaeobiological understanding of “Keratosa”-type non-spicular demosponge fossils in carbonates: with a new example from the lowermost Triassic, Armenia

**DOI:** 10.1101/2022.07.14.500008

**Authors:** Cui Luo, Yu Pei, Sylvain Richoz, Qijian Li, Joachim Reitner

## Abstract

Fossilised fibrous skeletons of non-spicular demosponges have been reported in carbonates throughout the Phanerozoic and recently from rocks dated back to 890 Ma ago. These records expanded our understanding of metazoans in multiple aspects, including their early evolution, the cryptic ecology in fossil reefs, and recovery after mass extinction events. However, debates and confusion persist on the differentiation of these fossils from other biogenic or abiotic structures. Here we propose six identification criteria based on established taphonomical models and biological characteristics of these sponges (“Keratosa” *sensu* Minchin, 1900). Besides, sponge fossils of this kind from the lowermost Triassic of Chanakhchi (Armenia) are described with a 3-D reconstruction to exemplify the application of these criteria in the recognition of such organisms. Subsequently, the state-of-the-art understanding of the taxonomy and evolution of these fossil sponges, a previously poorly addressed topic, is summarized. The morphology of the Triassic Chanakhchi fossils indicates an affinity with Verongimorphs, a group that may have evolved by the Cambrian Age 3. Other than that, more efforts are still required to explore the taxonomic diversity and evolutionary details in the accumulating data of these fossil non-spicular demosponges.

## 1. Introduction

The fossilised fibrous skeletal frame of non-spicular demosponges was first recognized in carbonates based on Triassic and Devonian examples in a study initially aimed to provide references for the search for Precambrian unmineralized ancestral animals [1]. Since then, fossils of this kind have been extensively reported [2,3]. Some of these records are especially valuable in revealing previously unexpected ecological and evolutionary facts about sponges: they were able to build microbialite-like bioconstructions [4–6], have bloomed in the aftermath of reef deterioration events in the middle Cambrian and earliest Triassic [7–11], and was recently found in carbonates of 890 Ma old [12], significantly preceding the estimated age of 720 Ma that the poriferan lineage emerged (e.g., [13]).

This group of non-spicular demosponges possess organic fibrous skeletons and conform to the definition of “Keratosa” in Minchin [14]. Living taxa with such morphological characteristics are now assigned to subclasses Keratosa and Verongimorpha based on phylogenomic studies [15–17]. Although it is difficult to study the exact taxonomy of these organisms in fossil materials due to the lack of histological, biochemical, and genomic information, the family Vauxiidae from Cambrian shale Lagerstätten has been assigned to the subclass Verongimorpha without many controversies because of its exceptionally preserved chitinous, cored skeletal fibres [18,19].

By contrast, let alone taxonomy, the recognition and interpretation of these non-spicular demosponge fossils in carbonates are still controversial, although differences between these fossils and superficially similar structures of other origins have been discussed in early literature [1,5]. Structures that seem to be most often confused with “Kearatosa”-type non-spicular demosponge fossils include lithistid sponges [20], Wedl tunnels [21], *Lithocodium* [22], amalgamated micritic clots [23], and metazoan burrows [24]. Since the identification of these non-spicular demosponge fossils in carbonates becomes increasingly influential to our understanding of metazoan evolution and paleoecology, it is necessary to re-examine and refine their recognition criteria. In addition, as data accumulates, the taxonomy, evolution, and other biological features of these organisms become further topics to be investigated.

This study tries to extract a set of recognition criteria of “Keratosa”-type non-spicular demosponge fossils in carbonates based on established taphonomic models and morphological characteristics of these organisms. A new fossil example from the lowermost Triassic carbonates is analysed to demonstrate the application of these criteria and methods. Finally, the state-of-the-art understanding of the taxonomy and early evolution of these non-spicular demosponge fossils are addressed.

## 2. Materials & Methods

As the basis of proposing identification criteria, the established taphonomic models of these organisms were first revisited. Then the six recognition criteria were proposed based on published data to avoid any circular argument in this article. They are generally a modification of the four morphological characteristics that Luo & Reitner [5] used to identify non-spicular demosponge fossils in thin sections. The information involved in the improvements can all be found elsewhere in the published descriptions in Luo & Reitner [1,5] and Luo [2].

To demonstrate the application of the proposed recognition criteria, a new example of “Keratosa”-type non-spicular demosponge fossil was analysed. The investigated materials were collected from the Chanakhchi (formerly also known as Zangakatun or Sovetashen) section in SW Armenia [8,25]. The section was located on the western margin of the Cimmerian microcontinent between the Neotethys and Palaeotethys oceans during the Early Triassic [26]. Carbonates in this section were deposited between fair weather and storm wave base on a distal and low-relief open-marine ramp [8,9]. Two levels of sponge-microbial bioherms are present above the end-Permian mass-extinction records. The lower one is 5 m thick, expanding from the post-extinction Permian to the second conodont zone of the Griesbachian, Induan (*Isarcicella isarcica* zone). The second one, late Griesbachian in age, is 13 m thick and encloses several thrombolites and dendrolitic biostromes and bioherms. The thickest microbialite deposit in the Chanakhchi section is from the upper microbialitic interval and is up to 8 m wide and 12 m thick [8]. Massive asymmetric thrombolitic domes characterize the lower half of the bioherm, followed by several thrombolitic biostromes and again several thrombolitic and dendrolitic mounds. The sample studied here comes from the lowest of the upper dendrolitic part and corresponds to the Sponge Facies 3 and sample 81 described in Friesenbichler et al. [8].

Thin sections were examined using a Zeiss SteREO Discovery.V8 microscope and photographed using the attached AxioCam MRc 5-megapixel camera. A chosen piece of the samples was cut into an approximately 30 × 30 × 5 mm chip for further 3-D reconstruction. It was mounted on a glass and then serially ground using the same method as that described in Luo & Reitner [1]. One hundred polished planes were photographed using the same set of microscope and camera system mentioned above. A Mitutoyo micrometre was used to control inter-plane distances. The average distance is 9.6 ± 1.1 μm (Supplementary File 1A). Although the 36^th^ and 37^th^ images were recorded the same by mistake, this does not much affect the outcome.

The obtained images were aligned using Adobe Photoshop and converted to the grey-scale mode (Supplementary File 1B). The 3.09 × 2.33 mm area illustrated in Fig. 4B, which includes a well-preserved skeletal frame and aquiferous canals, was cropped for further processing (Supplementary File 1C). The brightness and contrast of these images were adjusted one by one using GIMP 2.10.14. The dark areas in the background, which would otherwise affect the visualization of the skeletal frame, were meanwhile masked by a light grey colour (Supplementary File 1D). The resulted stack was visualized using Voreen 5.2.0 (voreen.uni-muenster.de) (Supplementary File 1D, 3A).

The smaller areas illustrated in Fig. 4C–D and 4G were processed similarly, but with the background more carefully removed in GIMP to better visualize the aquiferous canals and skeletal meshes (Supplementary File 1E–H).

All fossils and thin sections illustrated in this study are deposited at the University of Göttingen. Electronic data generated in the grinding and image processing are available in Supplementary File 1.

## 3. Results

### 3.1 Preservation of “Keratosa”-type demosponges in carbonates

Similar to the preservation of siliceous sponges (Fig. 1A-D), the fibrous skeleton of “Keratosa”-type non-spicular demosponges is often moulded in a micritic matrix and cemented by microspars (Fig. 1E). These structures are readily observable in thin sections but either discernible [1,5,11] or indiscernible [4,12] in polished slabs, depending on different materials. The organic skeletons of non-spicular demosponges are composed of spongin and/or chitin that are more resistant to biodegradation than other soft tissues [27–30]. This allows them to be moulded in syndepositional micrites and then replaced by calcite spars, the same as the taphonomic processes that siliceous spicules are subjected to.

**Figure 1.**
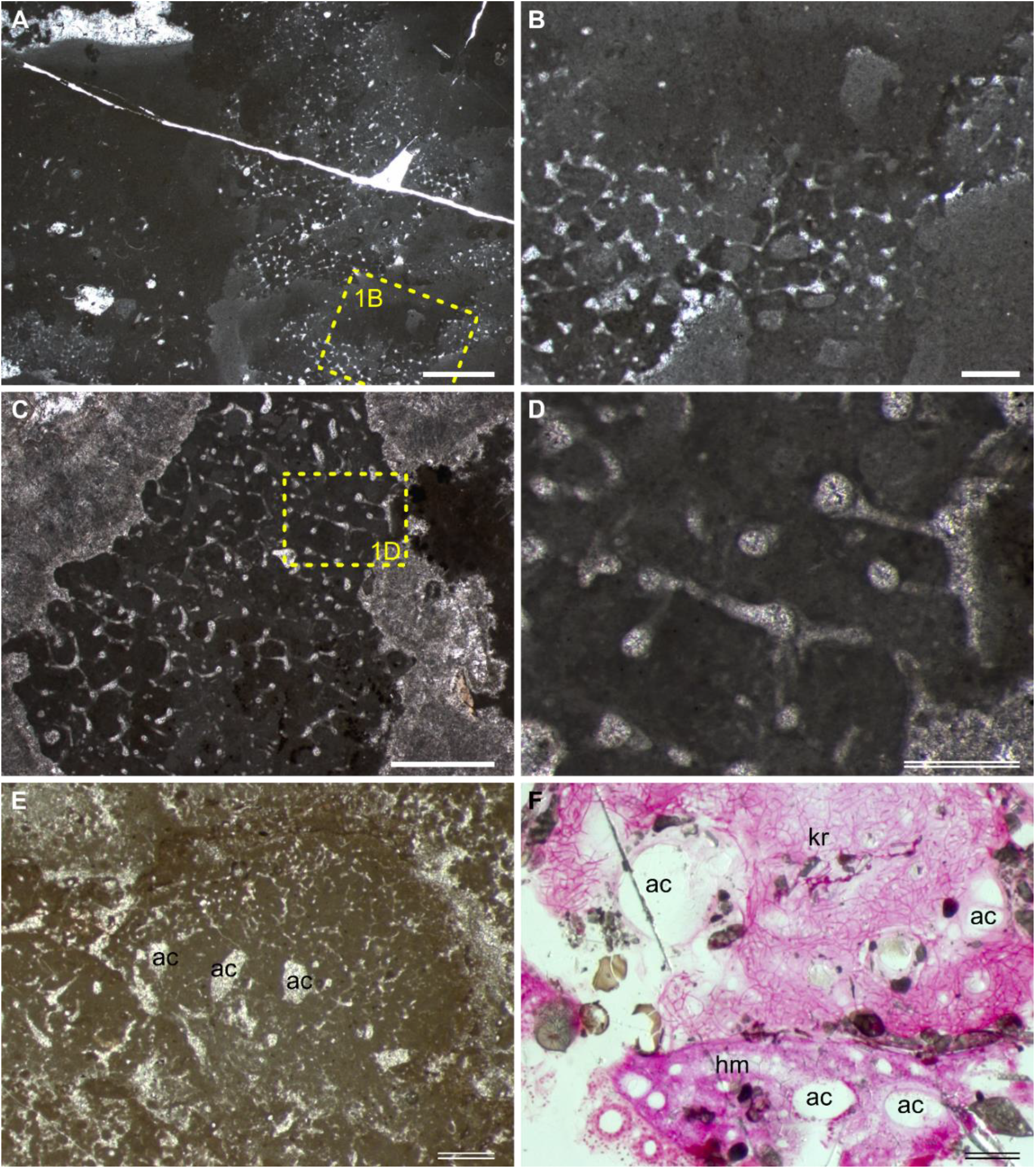
Preservation of sponges in carbonates. (A–D) A hexactinellid (A–B) and a lithistid (C–D) fossil from an Albian (Lower Cretaceous) mud mound in Araya, Spain. Rectangles in (A) and (C) are enlarged in (B) and (D), respectively. (E) A non-spicular demosponge fossil from the Carboniferous Clifton Down Limestone, UK. (F) Histological thin section of a living non-spicular demosponge (kr) and a homoscleromorph (hm) from the Lizard Island, stained in basic fuchsine. Comparable aquiferous canals (ac) are indicated in (E) and (F). Thin section number: (A–D), AR; (E), BH10; (F), Liz234. Scale bars: single line = 2 mm; double line = 0.5 mm.

The syndepositional micrites could be accumulated through two different paths. The first is the precipitation of automicrites during the decay of the sponge soft tissue. These processes and the resulting fossils have been repeatedly observed in modern and palaeontological examples [31–34]. The main trigger of the rapid automicrite formation has been attributed to the raised alkalinity and the presence of nucleation template due to the decay of sponge tissues in restricted microenvironments [32,35,36]. Some studies emphasize the role of organic sorbent in carbonate nucleation [34,37]. Generally, high alkalinity in the seawater is favourable for the automicrite precipitation.

The second path is the deposition of allomicrites. Take the hexactinellid fossil in Fig. 1A-B as an example. The skeletal frame seems to be first moulded by automicrites which show patched texture. Then the spongocoel was filled by allomicrites which form geopetal structures. The organic skeletons of non-spicular demosponges are for a longer time resistant against degradation and could be washed out from other soft tissues of the dead sponge (imagine a piece of natural bath sponge). Rapid burial of these skeletons with fine sediments would be favourable for their preservation.

### 3.2 Recognition criteria of “Keratosa”-type demosponges in carbonates

A perfect validation for the presence of these sponges would be an iconic sponge body with a spongocoel surrounded by an anastomosing fibrous skeletal frame. However, regardless of the fact that many shallow water and cave-dwelling non-spicular demosponges are encrusting and formless [38,39], a sponge body erected in the seawater is difficult to be completely fossilized according to the introduced taphonomic models. Almost all the “Keratosa”-type demosponges fossils so far described from carbonates are shapeless clots or layered encrusts, and many of them are only visible in thin sections.

For those fossils formed following the first taphonomic path, Luo & Reitner [5] used four morphological characteristics to identify them in thin sections and exclude other interpretations. These characteristics are here advocated again with a few refinements based on the information already provided in Luo & Reitner [1,5].

I. Fibrous skeletons are preserved as microspar-cemented moulds in homogeneous automicrites.
II. The skeletal fibres form an anastomosing network extending three-dimensionally in the micritic aggregation with a generally uniform density.
III. The skeletal fibres either persist in a uniform thickness or show variable thicknesses with regular orders or hierarchies. For reference, the diameters of skeletal fibres in living non-spicular demosponges vary from a few to hundreds of micrometres, with the majority being around tens of micrometres thick (Fig. 2; Supplementary File 2).
IV. The fibrous network is constrained in the micritic aggregation and exhibits fibres lining the border of the aggregation, e.g., between the sponge body and the hard substrates and wrapped particles.
V. There are no desmas (Fig. 1D), incorporated spicules, or orthogonal symmetry (Figs. 1A-B) in the fibrous network that could indicate an affinity of spicular sponges.
VI. Water canals of the sponge aquiferous system are sometimes preserved (Figs. 1E-F). If present, they add credits to the sponge interpretation.

**Figure 2.**
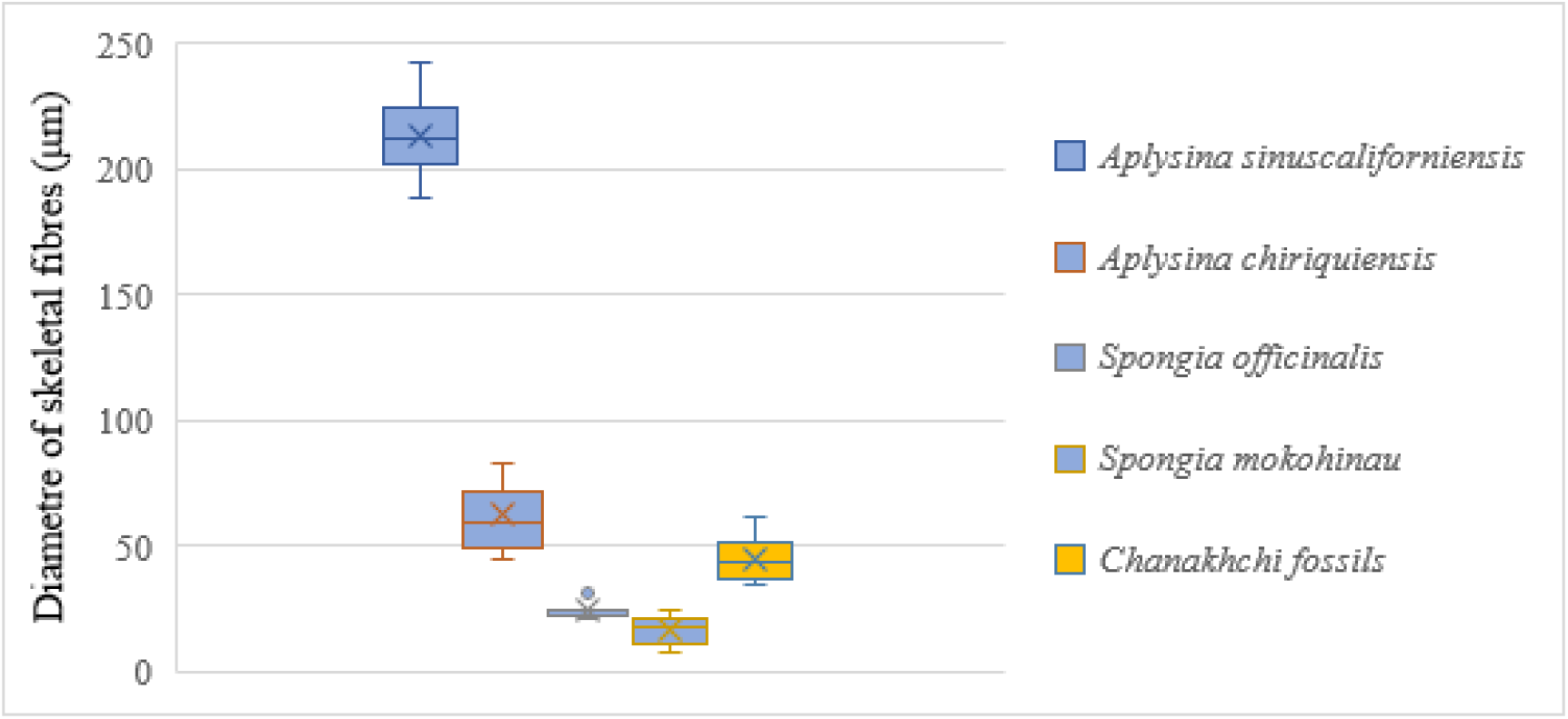
Thicknesses of skeletal fibres of a few living non-spicular demosponges compared with the Chanakhchi fossils. Original measurements and data sources see Supplementary File 2.

The above criteria II–III restrict the recognition of non-spicular demosponges to those groups that possess regular anastomosing skeletons, although many living taxa have dendritic skeletal frames and irregularly knotted fibres (e.g., family Pseudoceratinidae) [40]. However, to hold certainty on the recognition of these sponges in carbonate fossil records, this restriction is necessary at this stage.

For sponge skeletons preserved following the second path, many of these characters are inapplicable, including automicrites, the margin of the fibrous network, and aquiferous canals. The morphology of the 3-D fibrous network is the only thing that can be counted on. Preservation like this, as well as diagenetic alterations, could wipe away important biological information and introduce uncertainties to the identification of non-spicular demosponge fossils.

### 3.3 Observation of the Chanakhchi fossils

The thin sections contain bushy and rounded mesoclots, which are composed of microbially-induced crystal aggregates [8]. They grow on each other to form a dendrolitic column. Fossil sponges grew in the interspace of these thrombolites and dendrolites, sometimes with their upper surface preserved (Fig. 3A-B). No other microfossils have been observed in these materials.

**Figure 3.**
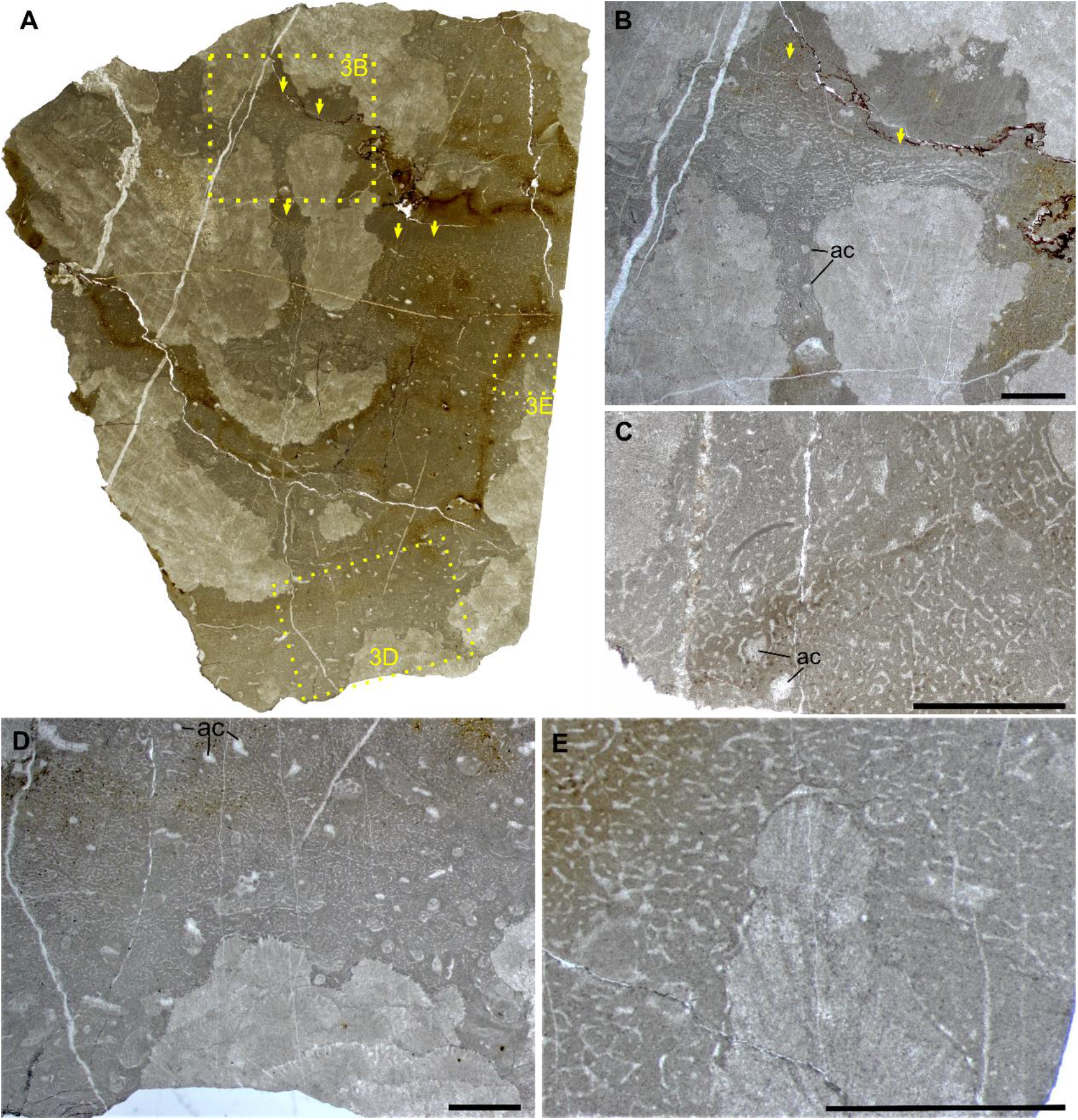
The sponge fossils from Chanakhchi (Armenia). (A) Overview of the sponge fossils encrusting over dendrolite. Areas in rectangles are enlarged in Fig. 3B, D, and E, respectively. (D–E) Close view of the skeletal fibres and aquiferous canals (ac). Yellow arrows in (A) and (B) indicate the upper surface of the organisms. Thin section number: (A–B, D–E), ZG; (C), ZG-small. All scale bars = 2 mm.

The sponge fossils are preserved as spar-cemented fibrous networks embedded in homogeneous grey micrites. The skeletal fibres are mainly 37–51 μm thick (Fig. 2; Supplementary File 2) and can line up the boundary between the organism and alien objects, such as the crystal fans and wrapped particles (Fig. 3B-E). No spicules and desmas have been observed in thin sections. Although the fibre thicknesses vary in a range, there was no clear separation of fibre hierarchies like that in many living dictyoceratid skeletons [40]. Serial grinding and 3-D reconstruction confirmed that these microspar-cemented fibres were indeed part of a three-dimensional network (Fig. 4; Supplementary File 3). Part of this network shows regular hexagonal meshes (Fig. 4E-G), although most meshes appear to be randomly polygonal. Many irregular but mostly tubular cavities are scattered in these fossils. They are one magnitude thicker than the skeletal fibres (200–300 μm wide in thin sections with extreme widths of over 500 μm, Supplementary File 2), cemented by calcite spars and sometimes containing geopetal fillings (Fig. 3B, D). These structures are different from fenestral fabrics in showing a rounded outline in cross-sections and lacking a layered distribution pattern. By 3-D reconstruction, these structures are proved blind-ending tubes, resembling the architecture of aquiferous canals in living sponges [41] (Fig. 4C-D). However, there are also rounded patches of the same scale as these cavities, which are, however, filled with skeletal fibres and paler micrites (Fig. 3D-E). These may represent either old aquiferous canals filled by a later generation of sponge tissues or different generations of micrite deposition during early diagenesis, similar to the paler patches in the spicular sponge fossils in Fig. 1B-C.

**Figure 4.**
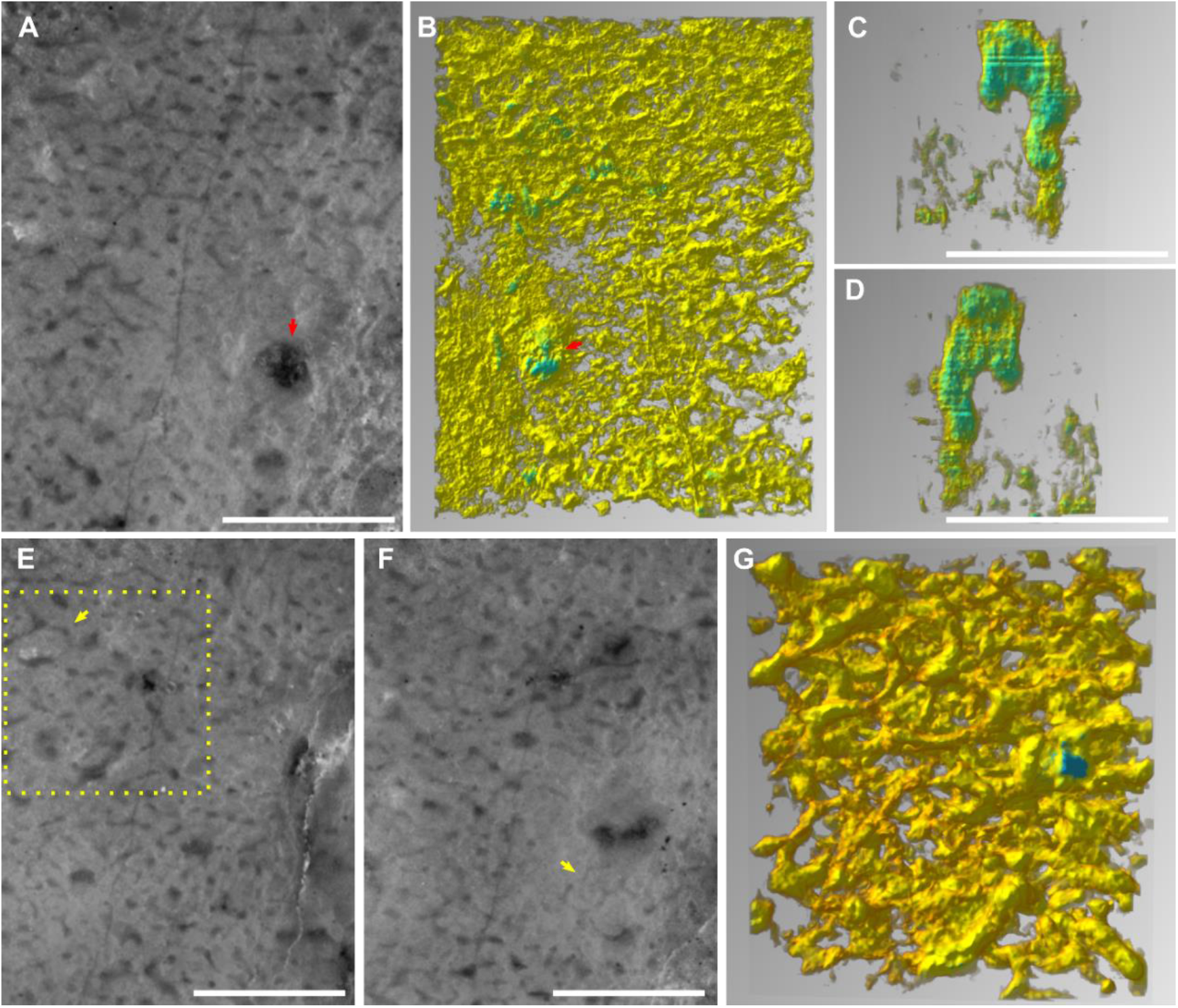
Serial grinding and 3-D reconstruction of a 3.1 × 2.3 × 1.0 mm chip of the Chanakhchi sponge fossil. (A–B) A serial grinding photo (A) and the corresponding 3-D reconstruction (B) show the presence of aquiferous canals (red arrows). The images of (A) and (B) mirror each other because (A) was the last plane of the serial grinding, and (B) shows the sole of the reconstructed block. (C–D) Lateral views of the isolated 3-D structure of the aquiferous canal, with the last serial griding plane on the top. (E–F) Serial grinding photos showing the existence of regular hexagonal meshes (yellow arrows). (G) 3-D reconstruction from the rectangle area in (E), showing the 3-D structure of the hexagonal meshes. All scale bars = 1 mm.

## 4. Discussion

### 4.1 Differentiating “Keratosa”-type demosponge fossils from other similar structures based on proposed criteria

As introduced in the beginning, several sorts of structures are easily confused with “Keratosa”-type demosponge fossils. Due to varied preservation types and quality, there is indeed a spectrum of morphological intermediateness between these demosponge fossils and structures of other origins. This holds true for nearly all sorts of fossil materials, and palaeontologists often cope with this problem by analysing only the best specimens. For this reason, when discussing the differentiation between “Keratosa”-type demosponge fossils and similar structures, we refer to the best specimens which exhibit all the proposed characteristics I–VI. Such fossil materials are not just a conception or fantasy. The Chanakhchi fossils have shown an example of them.

The morphological differences between the fibrous sponge skeletons and fossilized cyanobacteria and fungal hyphae have been discussed in Luo & Reitner [1,5]. Those arguments are still valid today. In addition, the above-stated characteristic-I can also help to discriminate. Fossilized filamentous cyanobacteria and fungal hyphae often have a dark lining along the fibres, representing their durable sheaths and cell walls, respectively [42,43]. In contrast, the moulds of sponge skeletal fibres do not have such a lining. The recognition of fossilized fungal hyphae networks is normally based on the presence of various reproductive structures like conidia or chlamydospores, which were not seen within the sponge skeletal fibres [44].

The morphological characteristics II–IV reflect the biological features of a sponge skeleton. The skeletal frame is a supportive structure and thus extends three-dimensionally with regularities in mesh size and shape and fibre thicknesses (characteristics II–III). Because it belongs to an individual animal, the fibrous network must have a clear border against the surroundings (characteristic IV). In contrast, Wedl tunnels, the labyrinth-like micrometre-scale canals, were bored by fungi or cyanobacteria to seek nutrients [45]. They do not have the geometric regularity and the self-constraining border as the sponge skeletons. *Lithocodium* is a problematicum possessing septate and branching filaments [46,47]. These filaments are not network forming, and their diameters decrease with successive branching. Compared with these non-skeletal structures, the skeleton of other sponges could be more easily confused with that of non-spicular demosponges [20]. In this case, characteristic-V could be consulted to discriminate these different skeletons.

It has been addressed previously that fenestral structures, compacted peloids, and amalgamated micritic clots do not form cavities with the regularity seen in sponge skeletons (characteristics II–IV) [1,5]. The Silurian structures described in Kershaw et al. [23] are similar to non-spicular demosponge fossils (e.g., Figure 3 in [23]) while also showing irregularities in other parts of those materials (e.g., Figure 1 in [23]). However, for the reason stated at the beginning of this section, the irregularities in this example are not a sufficient argument to invalidate the sponge interpretation in general. To establish amalgamated micritic clots as a general interpretation of those fibrous structures, it requires experiments or observations of modern examples in which compacted micritic clots do form uniform and complex cavity networks conforming to characteristics I–VI.

The Cambrian microburrows mentioned in Kris and McMenamin [24] were initially identified following an earlier description of similar structures in Wood et al. [48]. However, the diameter of these microburrows is 100–500 μm, much thicker than the fibre thicknesses of so far recognized fossil non-spicular demosponges [1,5,11](Fig. 2). There was no information about the 3-D morphology of these microburrows. The graphoglyptid trace fossils referred to in Kris and McMenamin [24], particularly *Paleodictyon*, typically occur in deep water siliciclastic sediments. They are composed of tunnels with millimetric to centimetric diameters, and the tunnels form regular hexagonal meshes mainly on two-dimensional surfaces parallel to the sea bed [49–51].

Despite the examples above, skeletons of non-spicular demosponges can still appear similar to many other structures in thin sections, such as plant roots moulded in caliche nodules [52], the trabecular meshwork in echinoderm skeletal plates [53], and some scleractinian corals in certain cross-sections [54]. Therefore, reliable recognition of non-spicular demosponges cannot be merely based on thin sections or thin section photos. All available information from outcrops to microfacies analyses must be considered synthetically to minimize ambiguity.

### 4.2 Identifying Chanakhchi fossils as non-spicular demosponges

According to the description in Section 3.3, the morphology of the Chanakhchi fossils clearly fits the identification criteria I–VI proposed in Section 3.2, and none of the alternative interpretations discussed in Section 4.1 could explain the combination of all these characteristics.

The development of the crystal aggregates in the studied materials indicates a carbonate-oversaturated setting. Therefore, the presence of the Chanakhchi sponge fossils fits the taphonomic model of “Keratosa”-type non-spicular demosponges in carbonates – micrites could be rapidly precipitated from the oversaturated seawater once the decay of the sponge tissue releases organic substrates for nucleation. Moreover, the preservation of the Chanakhchi material is comparable with that of the hexactinellid fossil illustrated in Fig 1A-B in showing a first automicrite and a second allomicrite generation.

### 4.3 “Keratosa” in the fossil record: known and unknown

Although occurrence data of “Keratosa”-type sponge fossils in carbonates are accumulating [3], most studies either only briefly mention the existence of these fossils or focus on their palaeoecological roles (e.g., [7,10,11,55]). The basic understanding of the diversity, taxonomy, and evolution of these fossil organisms has not yet been established.

The recently reported possible non-spicular demosponge fossils from the 890-Ma-old Little Dal Group [12] look nearly indistinguishable from the middle Cambrian to Ordovician analogues in thin sections. This raises a dilemma. On the one hand, there are yet no competitive alternative interpretations for these fossils other than non-spicular demosponges. On the other hand, if this interpretation is true, it’s hard to imagine how these organisms maintained their shape nearly unchanged through around 400 million years.

Regardless of this oldest example, perhaps the early Cambrian fossil record is the best starting point to explore the evolutionary history of these animals. After all, the most widely acknowledged fossil representatives of these demosponges are vauxiids from the Konservat-Lagerstätten of early to middle Cambrian shales [18,56–58]. Morphological characters of sponge skeletal frames can easily be obtained in this preservation art compared with that in carbonates. Vauxiids were assigned to the order Verongiida, subclass Verongimorpha, because their skeletons seem to be constructed with cored fibres composed of chitin [18,19], consistent with the skeletal characters of living verongiids [30,40,59]. Nevertheless, the newly described vauxiids from Cambrian Stage 3 and 5 of South China are all associated with silicification in the fibrous skeletons [56–58]. It requires further efforts to determine whether the silicification is of biological or diagenetic origin and whether this will affect the original phylogenetic assignment.

In records prior to the Cambrian Age 2, some fossils from the Tommotian carbonates of Siberia have once been interpreted as “Keratosa-”-type non-spicular demosponges (Fig. 5) [2]. These fossils were later diagnosed as the archaeocyath *Dictyocyathus translucidus* [60], a unique archaeocyath species whose skeleton is always preserved as moulds filled with sparitic cements [61–63]. This exceptionality in archaeocyaths was previously attributed to originally aragonitic mineralogy in the skeleton [61,62]. However, this interpretation does not explain the source of the micrites that often mould the spar-cemented skeletons. Published studies and illustrations of *D. translucidus* are very sparse. At least in the example illustrated in Fig. 5A, *D. translucidus* is preserved next to other archaeocyaths and *Epiphyton* shrubs that are devoid of micrite fillings in the inner- and interspaces. In this case, the non-spicular demosponge interpretation of *D. translucidus* seems to be reasonable from a taphonomic point of view.

**Figure 5.**
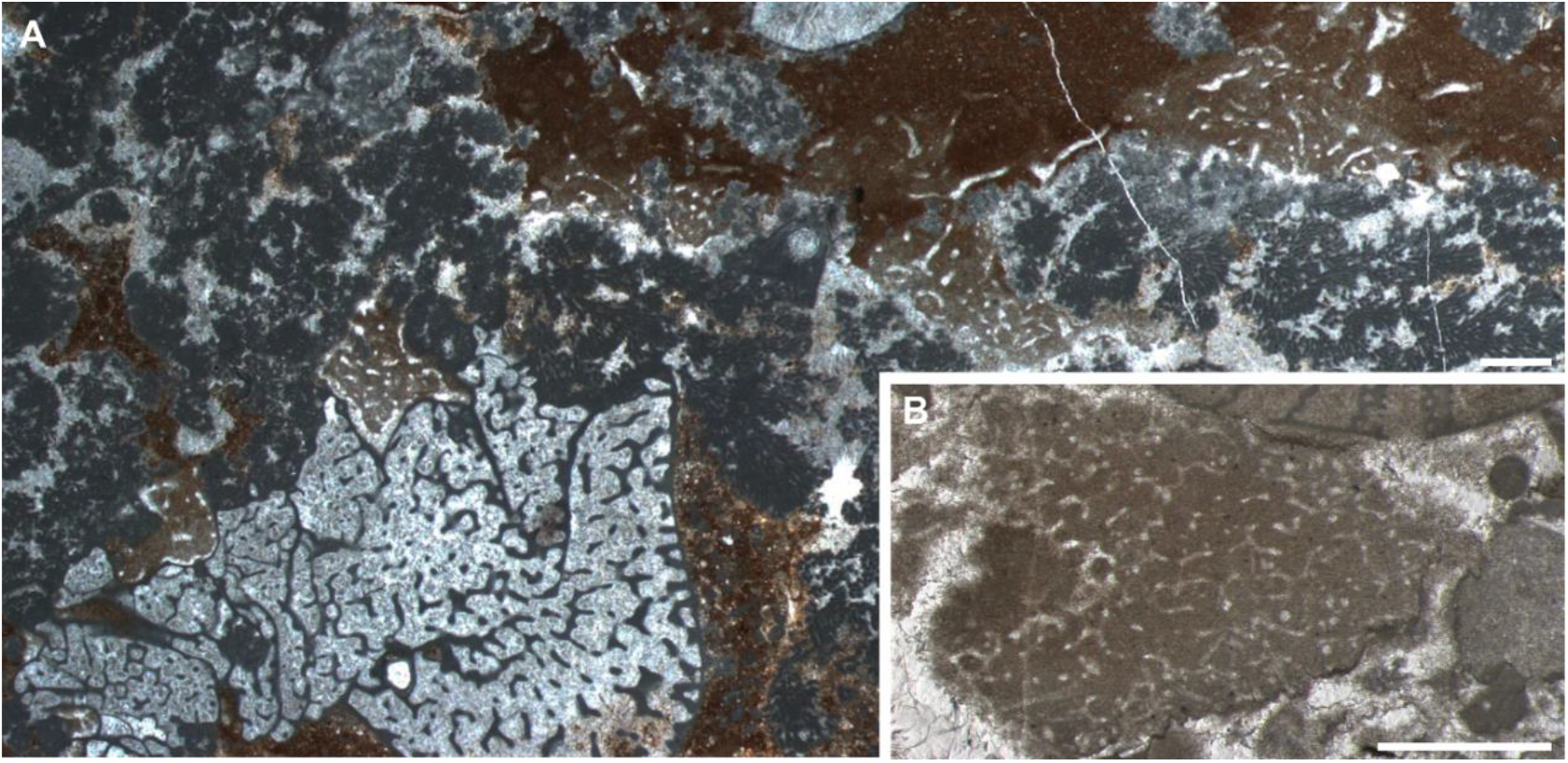
*Dictyocyathus translucidus* from the Tommotian archaeocyath-calcimicrobe reefs of Siberia. Thin section number: (A), CHD; (B), BB3. All scale bars = 1 mm.

Moreover, a recent study of fossils from the Guanshan Biota (Cambrian Stage 4) suggested that archaeocyaths of the suborder Archaeocyathina, to which *D. translucidus* also belong, are morphologically very close to vauxiid sponges [63]. The investigated Guanshan fossils can be comfortably assigned to either of these groups. The only problem is still the skeletal composition: the fibrous skeletons in the Guanshan fossils are silicified, neither carbonate as that of archaeocyaths nor organic as iconic vauxiids.

Noticeably, the skeletons of vauxiids and related archaeocyaths all show repeatedly occurring hexagonal-dominant meshes [63]. In living non-spicular demosponges, taxa that typically exhibit hexagonal meshes are verongimorphs such as *Aplysina* [64] and *Verongula* [65]. Although hexagonal meshes can also be sporadically present in the skeletal frame of some dictyoceratids such as *Spongia officinalis* [65], the Cambrian fossils do not show the hierarchical separation of fibres that is common in dictyoceratids [17,40,65]. Thus, these skeletal characters with non-hierarchical fibres and hexagonal meshes support the previous assignment of vauxiids to the order Verongiida. Following this comparison, the Chanakhchi sponges may also belong to Verongimorpha because their skeletons are dictyonal and without the separation of fibre hierarchies, although the frequency and regularity of hexagonal meshes in these fossils are incomparable with that of vauxiids.

Since subclasses Verongimorpha and Keratosa are sister groups, the order Dictyoceratida and part of order Dendroceratida are also expected to be present among the fossil specimens with reticulate fibrous skeletons. More morphological studies are required to explore further taxonomic and evolutionary details in these fossil materials.

## 5. Conclusion

By proposing the six identification criteria, commenting on alternative interpretations, and analysing a new fossil example, this study underpins the point that “Keratosa”-type non-spicular demosponges were indeed preserved in Phanerozoic carbonates. Recognizing these fossils requires a synthetical consideration of information from outcrops to lab analyses, from taphonomy to ecology, so that ambiguity and confusion can be minimized. Although such fossils are known throughout the Phanerozoic or even possibly since the Neoproterozoic, their taxonomy and evolutionary pattern are still poorly understood. However, if verongimorphs had evolved by the Cambrian Age 3 as discussed, some diversity is expected to be present in the fossil record. Comprehensive morphological studies combing with modern spongiology are required to reveal more details on this issue in the future.

## Supplementary Materials

Supplementary File 1 Raw data of the 3-D reconstruction. (A) Grinding records. (B) Originally photographed images aligned by Photoshop. (C) Cropped and grey-scale-adjusted images used for the reconstruction of Supplementary File 3A. (D) Files generated during the processing of Supplementary File 1C using GIMP and Voreen. (E) Cropped and grey-scale-adjusted images used for the reconstruction of Supplementary File 3B. (F) Files generated during the processing of Supplementary File 1E using GIMP and Voreen. (G) Cropped and grey-scale-adjusted images used for the reconstruction of Supplementary File 3C. (H) Files generated during the processing of Supplementary File 1G using GIMP and Voreen.

Supplementary File 2 (A) Measurements of the fibre thickness in some living non-spicular demosponges and the Chanakhchi fossil. (B) Images that were measured.

Supplementary File 3 (A) 3-D view of the region illustrated in Fig. 4A-B, E-F. (B) 3-D view of the aquiferous canal illustrated in Fig. 4C-D. (C) 3-D view of the skeletal frame illustrated in Fig. 4G.

## Acknowledgements

We appreciate A. Zhuravlev for the inspiring discussion, A. Hackmann and W. Dröse for helping with the serial grinding, and C. Hundertmark for the processing of serial grinding photos. We also had the logistic support of the National Academy of Sciences of Armenia, in particular Drs. Arkadi Karakhanyan and Lilit Sahakyan.

## Author Contributions

Conceptualization, C.L. and J.R.; Methodology, C.L. and J.R.; Investigation, C.L.; Resources, S.R. and J.R.; Data Curation, C.L. and J.R.; Writing – Original Draft Preparation, C.L.; Writing – Review & Editing, all authors; Visualization, C.L.; Supervision, J.R.; Funding Acquisition, C.L. and S.R.

## Institutional Review Board Statement

Not applicable

## Informed Consent Statement

Not applicable

## Data Availability Statement

The data presented in this study are openly available in FigShare at https://doi.org/10.6084/m9.figshare.c.6089724.v1.

